# Integrative genomic, transcriptomic, and epigenomic analyses of benign prostatic hyperplasia reveal new options for therapy

**DOI:** 10.1101/805168

**Authors:** Deli Liu, Jonathan Shoag, Daniel Poliak, Ramy S. Goueli, Vaishali Ravikumar, David Redmond, Aram Vosoughi, Jacqueline Fontugne, Heng Pan, Daniel Lee, Domonique Thomas, Keyan Salari, Zongwei Wang, Alessandro Romanel, Alexis Te, Richard Lee, Bilal Chughtai, Aria F. Olumi, Juan Miguel Mosquera, Francesca Demichelis, Olivier Elemento, Mark A. Rubin, Andrea Sboner, Christopher E. Barbieri

**Affiliations:** Sandra and Edward Meyer Cancer Center, Weill Cornell Medicine, New York, NY, USA; Department of Urology, Weill Cornell Medicine, New York, NY, USA; HRH Prince Alwaleed Bin Talal Bin Abdulaziz Alsaud Institute for Computational Biomedicine, Weill Cornell Medical College, New York, NY, USA; Department of Radiology, Weill Cornell Medicine, New York, NY, USA; Englander Institute for Precision Medicine of Weill Cornell Medicine and NewYork-Presbyterian Hospital, New York, NY, USA; Department of Pathology and Laboratory Medicine, Weill Cornell Medical College, New York, NY, USA; Beth Israel Deaconess Medical Center, Harvard Medical School, Boston, MA, USA; Centre for Integrative Biology, University of Trento, Via Sommarive 9, 38123, Trento Italy; Department of BioMedical Research, University of Bern and Inselspital, 3008 Bern, Switzerland

## Abstract

Benign prostatic hyperplasia (BPH), a nonmalignant enlargement of the prostate, is one of the most common diseases affecting aging men, but the underlying molecular features of BPH remain poorly understood, and therapeutic options are limited. Here we employed a comprehensive molecular investigation of BPH, including genomic, transcriptomic and epigenetic profiling of 18 BPH cases. At the molecular level, we found no evidence of neoplastic features in BPH: no evidence of driver genomic alterations, including low coding mutation rates, mutational signatures consistent with aging tissues, minimal copy number alterations, and no genomic rearrangements. Similarly at the epigenetic level, we found global hypermethylation was the dominant process (unlike most neoplastic processes). By integrating transcriptional and methylation signatures, we identified two BPH subgroups with distinct clinical features and associated signaling pathways, which were validated in two independent cohorts. Finally, our analyses nominated *mTOR* inhibitors as a potential subtype-specific therapeutic option. Supporting this, a cohort of men exposed to *mTOR* inhibitors showed a significant decrease in prostate size. Our results demonstrate that BPH consists of distinct molecular subgroups, with potential for subtype-specific precision therapy via *mTOR* inhibition.

## Introduction

Benign prostatic hyperplasia (BPH) is a common disease, affecting nearly all men as they age^1, 2, 3, 4, 5^. BPH frequently results in bladder outlet obstruction with concomitant lower urinary tract symptoms or infections, and more rarely bladder decompensation and renal failure^3, 6, 7^. The prevalence of BPH increases with age, with BPH symptoms reported by roughly half of men at age 50^1, 2, 3, 4, 7^. Approved medical therapies for BPH are limited to alpha-blockers, 5-alpha reductase inhibitors, and *PDE5* inhibitors^8, 9, 10^. However, many patients fail medical therapies, and require surgical intervention^11^. Histologically, BPH is characterized as the overgrowth of stromal and epithelial cells, and it occurs in the transitional zone of prostate^1^. Currently, many BPH studies have focused on risk factors of BPH^12, 13, 14, 15^, while the underlying molecular features of BPH remain understudied^3, 9, 16, 17, 18, 19^ and molecular data is relatively scarce^20, 21^. Moreover, BPH has been described as “the most common benign tumor in men”, and is commonly referred to as an adenoma, but unlike many malignant^22, 23^ and benign neoplasms^24, 25, 26^, it is unknown whether BPH is a neoplastic process ^3, 7, 18, 19, 20^. In this study, we performed a comprehensive investigation of 18 BPH cases via next-generation sequencing technology (Table S1). We selected samples from patients with very large prostates (top percentile and greater than 100cc, Table S1 and Figure 1A), based on the rationale that these “extreme outlier” were more likely to harbor biologically informative events^27, 28^.

**Figure 1.**
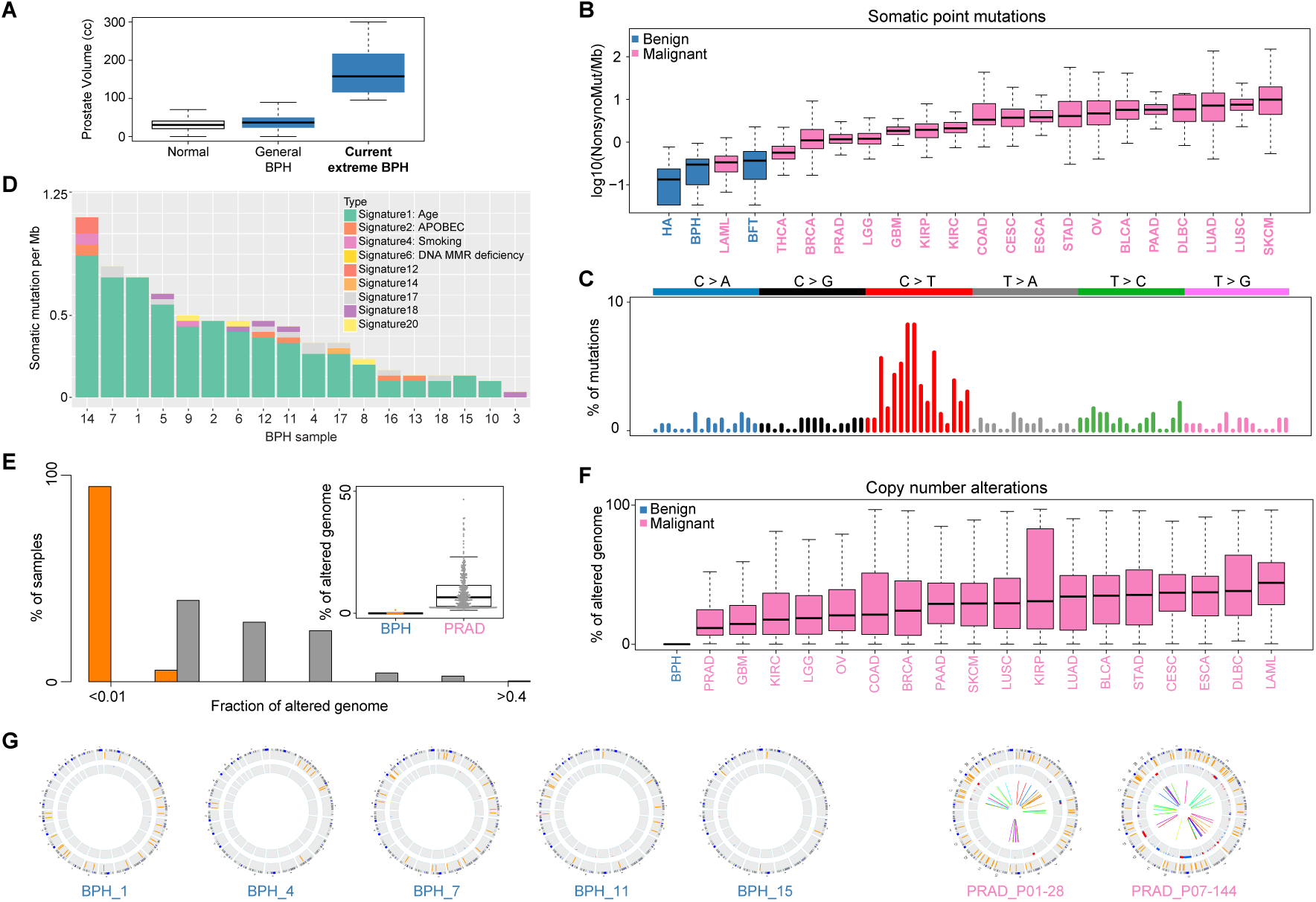
The low genomic alterations found in BPH samples. (A) Boxplots of prostate volume (cc) of normal, general BPH, and extreme BPH cases used in the current study. (B) The prevalence of somatic non-synonymous mutations across benign disease and multiple cancer types. The y-axis represents the log10 value of mutations. The x-axis includes benign (blue) and malignant tumors (pink) from TCGA studies. HA: hepatocellular adenomas, and BFT: breast fibroadenomas. (C) The somatic mutation signatures of BPH. The signature is based on the 96 substitutions classification defined by the substitution class and sequence context immediately 3’ and 5’ to the mutation position. The y-axis represents the percentage of mutations attributed to a specific mutation type. The six types of substitutions are shown in different colors. (D) The contribution of mutation signatures to each BPH sample. Each bar represents a BPH case and y-axis denotes the number of somatic mutations per megabase. (E) The fraction of altered genome, partitioned into bins covering a range from <0.01 to ≥0.4, shown as a histogram for BPH and primary prostate cancer samples. Inset: boxplot of altered genome fraction for BPH samples and primary prostate cancer samples from TCGA study. (F) The lower fraction of altered genome in BPH (blue) when compared to malignant diseases (pink) from TCGA studies. (G) Circos plots of 5 BPH and 2 primary prostate cancer samples. The rings from outer to inner represent somatic coding mutations, copy number alterations and genomic rearrangements respectively.

## Results

To define the landscape of genomic alterations in BPH, we performed whole genome sequencing (WGS), whole exome sequencing (WES) and SNP arrays on 18 BPH cases and matched controls (Table S1). The number of somatic coding mutations (SNV) ranged from 0.1 to 1 per megabase (Mb) (Table S2). As compared to neoplastic diseases (benign and malignant)^24, 25, 26^, BPH samples harbored fewer SNVs (Figure 1B), and there were no recurrent SNVs to suggest driver alterations. To understand underlying mutational processes, we examined mutational signatures^29^ across all BPH cases, and found BPH was highly associated with mutation signature 1^29^, which included C>T substitutions at NpCpG trinucleotides (Figures 1C and 1D). This signature has been shown to correlate with age^29, 30^, consistent with the age-related onset of BPH^1, 2, 3, 4, 7^. Moreover, BPH samples harbored minimal copy number alterations, and the fraction of altered genome was far lower than seen in primary prostate cancer^23^ and other neoplastic diseases (Figures 1E and 1F, Table S4 and S5). Also unlike primary prostate cancer^23, 31^, analyses of structural variants in WGS revealed no genomic rearrangements in BPH (Figures 1G and S1). Together, these data show no evidence of driver genomic alterations in BPH, inconsistent with a neoplastic disease process.

We next examined the transcriptional landscape of BPH using RNA-seq. Because BPH, by its very nature has no “adjacent normal” tissue, we compared the gene expression profiles from BPH samples with histologically normal transition zone tissue sampled from age-matched controls (Figure 2A). We identified a BPH transcriptional signature that included 392 differentially expressed genes between BPH and control samples (Figure 2B and Table S6). When compared to control samples from the normal peripheral zone, this transcriptional signature was BPH specific, and not specific to transition zone tissue (Figure S2). We next validated this BPH transcriptional signature using two independent study cohorts^21, 32^, and again found reliable clustering of BPH samples (Figures 2C and 2D) with similar upregulation of *BMP5* identified (Table S6). Having defined and validated a robust set of genes altered in BPH, we explored the signaling pathways deregulated using gene set enrichment analysis (GSEA)^33^ (Figure 2E). Interestingly, multiple signatures related to *inactivation* of *KRAS* signaling were observed in our dataset, with concordance in an independent cohort (Figure 2E), and again inconsistent with a neoplastic process. In addition, we observed AR signaling downregulated in BPH (Figure 2F), consistent with previous findings that AR signaling disruption correlated with prostate inflammation and BPH pathogenesis^34, 35^.

**Figure 2.**
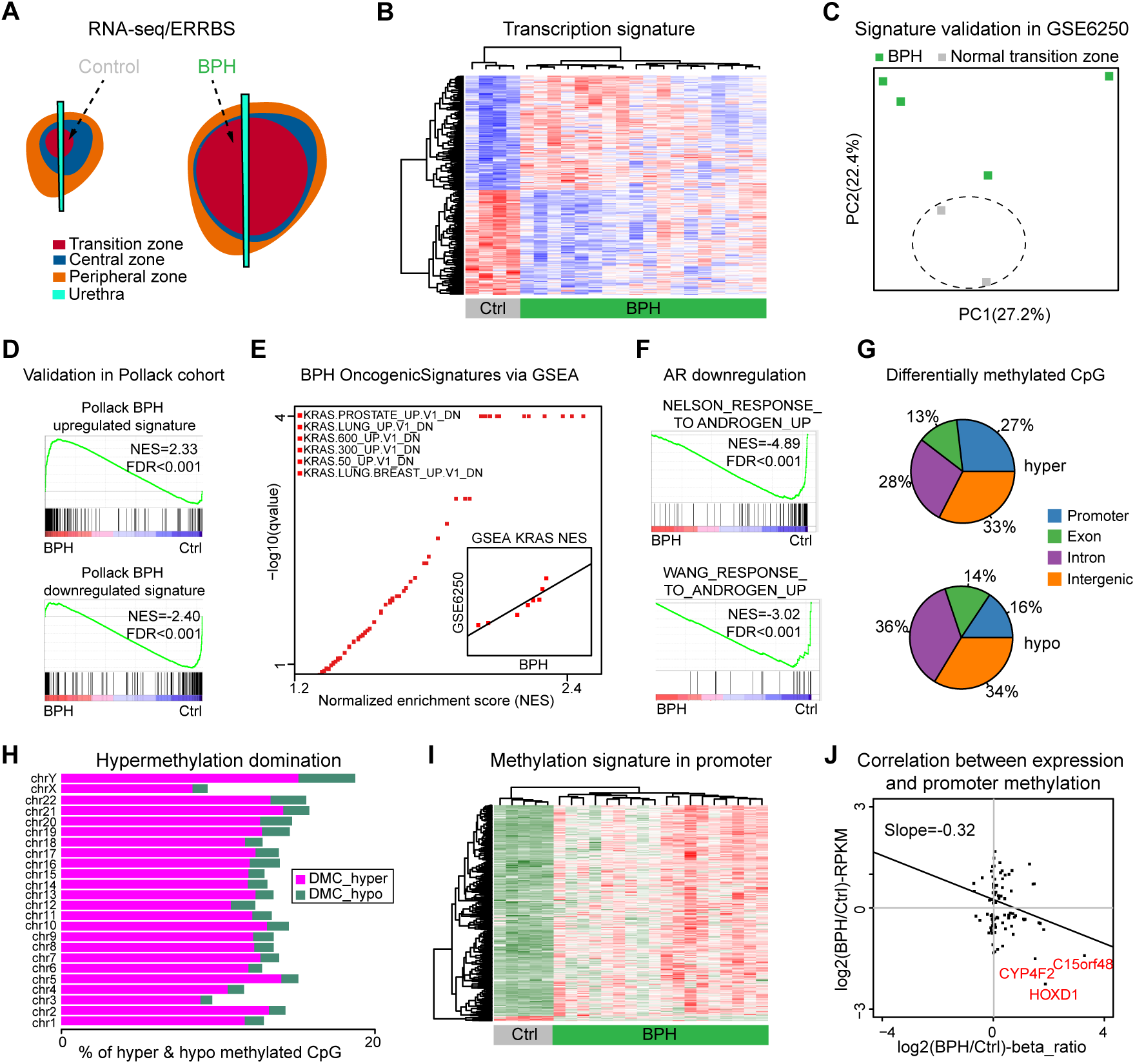
BPH transcription and methylation profiles. (A) Diagram of sampling location of BPH and control samples used for RNA-seq and ERRBS. Green color represents BPH samples, and grey color represents control samples from normal transitional zones. (B) Hierarchical clustering and heatmap of transcriptional signatures based on significantly differentially expressed genes between BPH and control samples. (C) Validation of transcriptional signature (panel B) in an independent study (GSE6250). Green color represents BPH samples, and grey color represents control samples from normal transitional zones. (D)The concordance of transcriptional signatures between current and previous BPH study. GSEA analyses of current BPH cases showing that genes upregulated in previous BPH cases are positively enriched, and genes downregulated in previous BPH cases are negatively enriched. (E) GSEA analysis of BPH cases in oncogenic signatures showing that genes downregulated in many cell lines when over-expressing an oncogenic form of *KRAS* gene are positively enriched. Similar results and high correlation with GSE6250 study are shown in inner panel. (F) GSEA analysis of BPH cases in AR related signatures showing that genes upregulated in LNCaP cells treated with synthetic androgen are negatively enriched. (G) Pie chart of differentially methylated CpGs between BPH and control samples among different genomic regions. Colors denote different genomic related regions. (H) Hypermethylation domination found in each of chromosome. The x-axis represents the percentages of hypermethylated (label as pink) and hypomethylated (label as green) CpGs. (I) Hierarchical clustering and heatmap of promoter methylation signature between BPH and control samples. (J) Correlation between transcription and promoter methylation signatures, and the examples of epigenetically silent genes are shown in red.

Next, we investigated the epigenetic landscape of BPH by defining the DNA methylation profile of 18 BPH samples and 5 controls from normal transition zone tissue using ERRBS (Enhanced Reduced Representation Bisulfite Sequencing). We identified 92,046 hypermethylated CpGs and 10,117 hypomethylated CpGs across different genomic regions in BPH, with hypermethylation being the dominant signal across all genomic regions, even when controlling for bias of CpG-rich loci (Figures 2G, 2H and S3). We defined a methylation signature for BPH that included 696 differentially methylated CpGs in promoter regions (Figure 2I and Table S7). Consistent with DNA methylation as a major mechanism of transcriptional control in BPH, we found negative correlation between promoter methylation and gene expression (Figure 2J). For instance, *HOXD1* was both underexpressed and hypermethylated at the promoter in BPH specimens, consistent with the downregulation of AR signaling pathway found in BPH^36, 37^ (Figures 2E and S3).

Identifying distinct molecular subtypes in human disease has provided insight into important biological and clinical phenomena. We therefore performed integrative analysis using transcriptional and methylation profiling, and identified two distinct BPH subtypes (Figure 3A and Tables S6-7), supporting robust biologically distinct subgroups across different data types. To validate distinct subtypes in BPH, we tested our signature via k-means clustering in two independent cohorts^20, 21^, and identified nearly identical subgroups (Figures 3C, 3E and Table S8), further supporting the robustness of these subgroups across data types and sources. We then examined the molecular and clinical features of these two groups. One subgroup (BPH-A) was enriched in stromal signatures^38^ (Figures 3B and S4), again in the validation cohort as well (Figure 3D and S5). Integrating the stromal cell signatures from single cell RNA-seq on normal prostate tissue^39^ further confirmed the stromal enrichment in BPH-A subgroup (Figure S7). Of note, there was no clear enrichment of stromal cell content visible on histopathology analysis of these samples, suggesting that molecular characterization provided independent information (Figure S8).

**Figure 3.**
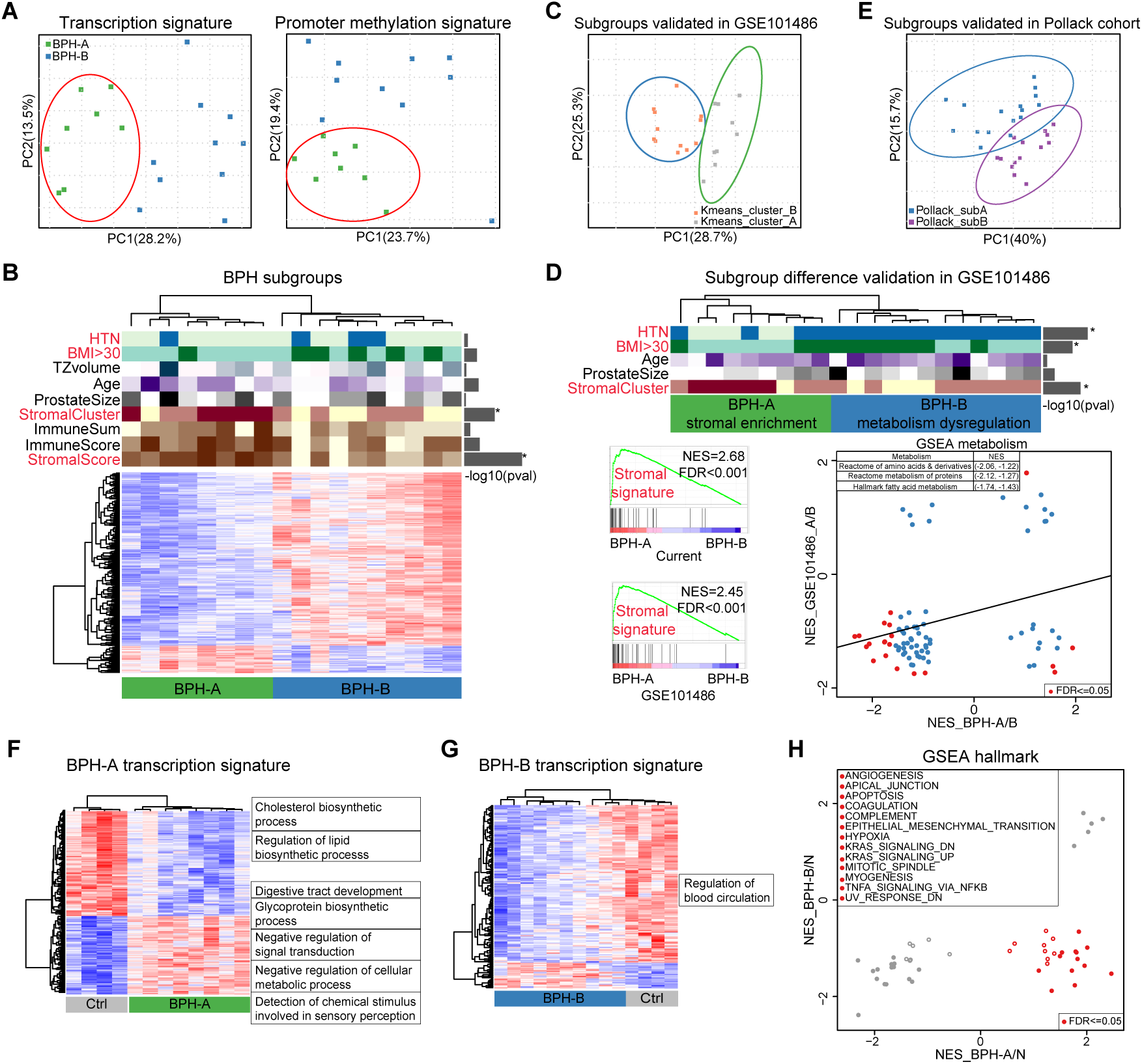
Identification and validation of distinct BPH subgroups. (A) Principal component analysis (PCA) based on transcriptional and promoter methylation signatures on RNA-seq data. Green denotes subgroup A, and blue denotes subgroup B. (B) The clinical and biological differences between two BPH subgroups. * denotes p-<0.05 assessing differences between two BPH subgroups. (C) The validation of BPH subgroups on an independent microarray study GSE101486 with 21 BPH samples via principal component analysis (PCA). K means clustering identified two distinct subgroups based on BPH subgroup signature from panel B. (D) Clinical and biological differences are shown between two subgroups from GSE101486 study. Bottom left represents GSEA plots of significant enrichment of stromal signature in subgroup BPH-A when comparing with subgroup BPH-B from both current and GSE101486 studies. Bottom right showed the correlation of metabolism dysregulation between two subgroups. X-axis denotes the normalized enrichment scores from current study, and y-axis denotes the normalized enrichment scores from GSE101486 study. Red dots represent the significant signature with FDR <0.05 in either one of two studies. Examples of metabolism dysregulation examples are shown. * denotes p-value <0.05 assessing differences between two BPH subgroups. (E) The validation of BPH subgroups on an independent RNA-seq study^21^ with 30 BPH samples via principal component analysis (PCA), based on BPH subgroup signature from panel B. (F) Hierarchical clustering and heatmap of transcriptional signature between subgroup BPH-A and control samples. The enriched functions are shown. (G) Hierarchical clustering and heatmap of transcriptional signature between subgroup BPH-B and control samples. The enriched functions are shown. (H) The difference of enriched pathways between BPH subgroups when comparing with control samples. Red dots indicate the difference of MSigDB hallmark signatures via GSEA with FDR ≤0.05 between two BPH subgroups. The x and y-axes represent the normalized enrichment score of signatures from each BPH subgroup when comparing to control samples.

The second subgroup (BPH-B) was enriched for patients with obesity (BMI >30) and hypertension (Figure 3D), potentially suggesting distinct pathobiology. Consistent with this, gene set enrichment analysis between the two subgroups demonstrated significant differences among metabolism related signatures, such as fatty acid and amino acid metabolism (Figure 3D). Positive correlation of metabolism dysregulation between the two subgroups extended to both cohorts (Figures 3D and S6), consistent with the clinical associations with obesity and hypertension. We then explored signaling pathways within each subgroup to further understand the underlying biology. As compared to control samples, we found differential expression of metabolism related genes predominantly in BPH-A samples (Figures 3F and 3G, Tables S10 and S11), consistent with the metabolism difference between two subgroups. Unbiased gene set enrichment analysis (GSEA) showed multiple deregulated pathways for each subgroup, with many pathways were negatively correlated between two subgroups (Figure 3H and Table S12), reinforcing distinct biology. Together these molecular data suggested two distinct biological categories of BPH – one with stromal-like molecular features, and the other associated with deregulation of metabolic pathways that presents in patients with underlying metabolic disturbances.

To nominate potential subtype specific therapeutic options, we utilized the Connectivity Map^40, 41^ analysis (Figure 4A and Table S13), which uses transcriptional expression data to probe relationships between diseases, cell physiology, and therapeutics. Strikingly, we found 50% of nominated compounds in BPH-A subgroup were related to inhibition of *mTOR* signaling (Figure 4B), and the subgroup enrichment of *mTOR* signaling was validated in two independent cohorts (Figure 4C), consistent with prior isolated reports in model systems^42, 43, 44^. To interrogate the potential effect of *mTOR* treatment on the prostate, we examined prostate size on cross-sectional imaging in patients taking *mTOR* inhibitors. We identified 425 male patients who had been prescribed an *mTOR* inhibitor (everolimus, sirolimus, or temsirolimus) for transplant or treatment of a non-prostate malignancy. We then reviewed these patient’s charts to identify men with accessible CT imaging including the pelvis before and after treatment, identifying 47 such subjects. CT scans from these 47 subjects and 12 men with serial imaging for nephrolithiasis (negative controls) then underwent blinded review and assessment of prostate size (Figure S9). Of these men, 17/47 had a prostate size decrease based on pre-established thresholds (12.5% decrease from baseline), all of whom were on an *mTOR* inhibitor (Figures 4D and 4F). None of the nephrolithiasis patients showed a significantly prostate size decrease. A higher proportion of patients taking Everolimus had a decrease in prostate size (pvalue=0.02) than Sirolimus (pvalue =0.06), compared to kidney stone patients (0%) (Figure 4E). Similar trends were seen in the effect of *mTOR* inhibitors on absolute cross-sectional area (Figure S10). Overall, these data suggest that subgroup BPH-A represents a biologically distinct subtype of disease preferentially dependent on *mTOR* signaling, and *mTOR* inhibition could serve as a novel therapeutic option.

**Figure 4.**
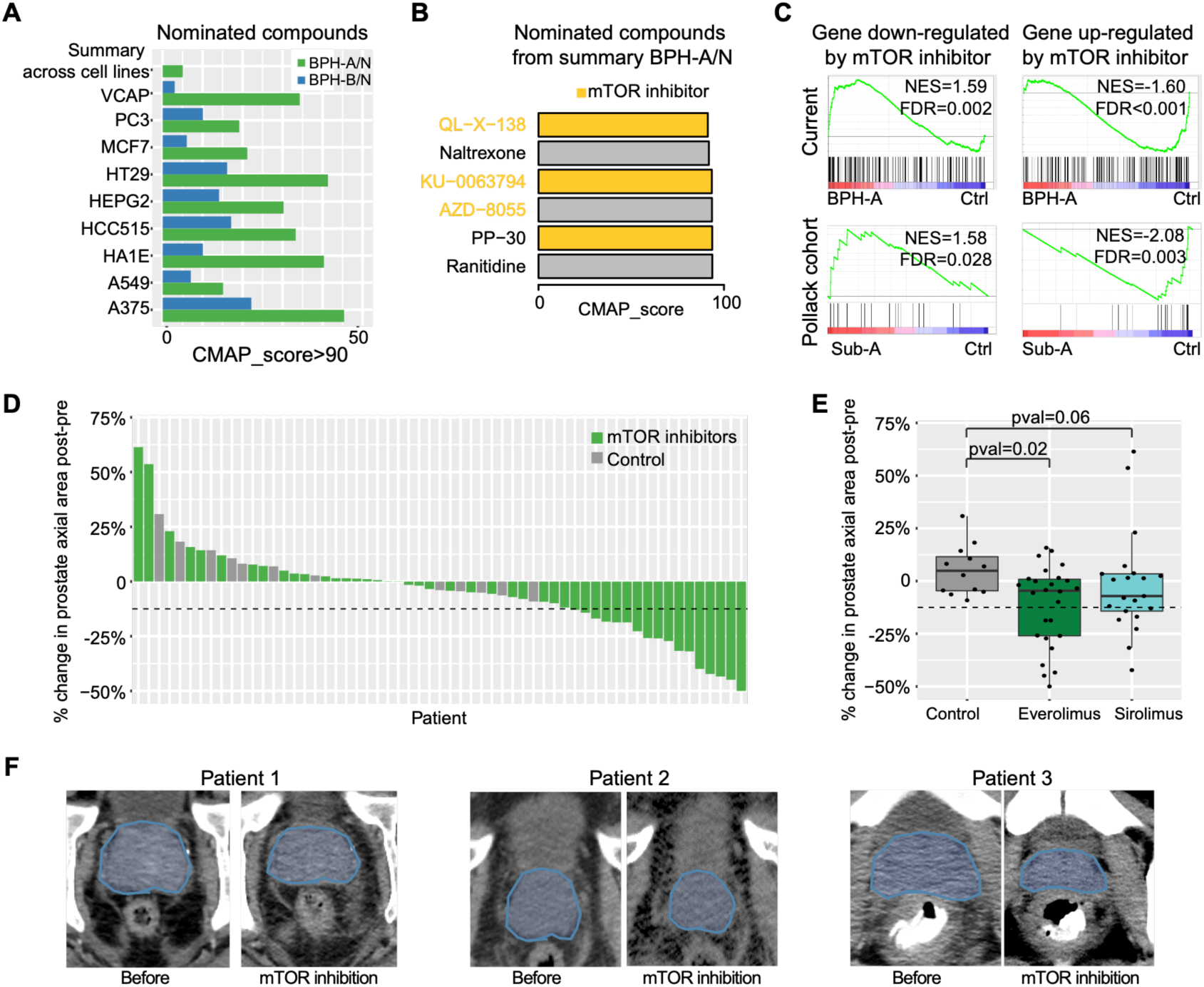
The alterations and potential precision therapies of BPH subgroups. (A) Barplots of nominated compounds from each BPH subgroup when comparing with control samples across multiple cell lines, and summary from all cell lines via Connectivity Map (CMAP). X-axis denotes CMAP score. Different colors represent BPH subgroups. (B) Nominated compounds from subgroup BPH-A via Connectivity Map. X-axis denotes the CMAP score. (C) GSEA analysis of *mTOR* related signatures in subgroup BPH-A and subgroup Sub-A from independent study^21^, showing that genes down-regulated by *mTOR* inhibitor are positively enriched, and genes up-regulated in CEM-C1 cells (T-CLL) by *mTOR* inhibitor are negatively enriched, when comparing to control samples. (D) Waterfall plot of % prostate axial area change on computed tomography in 47 patients after initiating therapy with an *mTOR* inhibitor and 12 kidney stone patients (negative controls). Different colors represent patient type. Dashed line represents a predetermined threshold (12.5%) for a significant decrease in area. (E) Boxplots of % prostate area change in 47 patients taking *mTOR* inhibitors (26 with Everolimus and 21 with Sirolimus treatments), and 12 negative controls. Pvalues represent prostate size change for each drug as compared to controls. (F) Examples of CT scans from three patients who had a decrease in prostate size after initiation of an *mTOR* inhibitor. Prostate highlighted in blue.

## Discussion

In summary, we report the first comprehensive, multi-level molecular investigation of BPH, including genomic, transcriptomic and epigenomic profiling. While dogma often suggests BPH represents a benign neoplastic process, we find no evidence of somatic genomic alterations, unlike benign neoplasms like such as frequent *MED12* mutations in breast fibroadenomas^24, 25^ and uterine fibromas^45^ or *FRK* mutations in hepatocellular adenomas^26^, and BPH exhibited an age-related mutation signature, consistent with the higher prevalence in older patients as opposed to underlying oncogenic processes. Furthermore, unlike the global hypomethylation signature in neoplastic diseases^46, 47, 48^, the DNA methylation landscape in BPH was dominated by hypermethylation. Together, our genomic and epigenomic data argues against BPH arising from a neoplastic disease process.

By integrating the transcriptional and DNA methylation data, we identified and validated two molecular subgroups in BPH, one characterized by a stromal signal (despite no clear differences in histology), and the other associated with hypertension and obesity, which was consistent with metabolism dysregulation between these two subgroups. Moreover, the altered signaling pathways of each subgroup comparing with control samples were related to the metabolism regulation and hypertension. Finally, we nominated potential therapeutic compounds for each BPH subgroup, and found that *mTOR* inhibitors may be preferentially active in one subgroup. By validating *mTOR* treatment on our institutional patient cohort, we found 17/47 patients treated with *mTOR* inhibitors showed the significant decreases on prostate size based on CT scan imaging, and no decrease found in patients without *mTOR* inhibitor. Overall, our findings provide critical insight into the underlying pathobiology of BPH, identify for the first time distinct molecular subgroups, and introduce a paradigm of precision therapy for a disease that affects the majority of aging men.

## Methods

### Samples collection

Patients with BPH (benign prostatic hyperplasia) were prospectively enrolled for sequencing of prostate tissue samples from transitional zones under a protocol approved by the institutional review board of Weill Cornell Medical College. Normal controls for RNA-seq and ERRBS were obtained from men undergoing radical prostatectomy for prostate cancer without BPH. Under the supervision of study pathologists, benign areas of transition zone distant from the tumor were cored. Written informed consent was obtained, including discussion of risks associated with germline sequencing. Fresh tissue samples were collected and processed using internal standard operating procedures.

### Whole genome sequencing and whole exome sequencing platform, data processing and analysis pipeline

Whole genome sequencing (WGS) on 5 BPH samples with matched normal samples from blood tissue was performed in New York Genome Center under standard protocol and pipeline for 100bp paired-end sequencing. Samples were sequenced with average genome coverage of 100x for BPH samples, and 50x for matched control samples. Whole exome sequencing (WES) on 13 BPH samples with matched control samples from blood tissue was performed in the Genomics Core of Weill Cornell Medicine under standard protocol and pipeline for 75bp paired-end sequencing. Whole exome sequencing capture libraries were constructed from BPH and control tissue by using SureSelected Exome bait (Agilent), and samples were sequenced with average target exon coverage of 300-360x for BPH and matched control samples.

Paired-end sequence reads of WGS and WES data were aligned to the human reference genome (hg19) using BWA^49^ v0.7.12. Sorted bam files were generate via SAMtools^49^ v.0.1.19, and the duplicated mapped reads were marked with Picard v1.134. BAM files were locally realigned to the human reference genome using GATK^50^ v3.7, and somatic base substitutions and small indels were detected by using MuTect^51^ v1.1.4 and Varscan2^52^ v2.3.9, with the sequencing coverage cutoff of at least 14x in BPH and 8x in control samples. Mutations were defined as the shared output between MuTect and Varscan2. After excluding the known human SNPs (dbSNP Build 150) and SNPs detected from control samples, the remaining mutations were annotated by ANNOVAR^53^ v2018.04.16 with GENCODE human gene annotation. The mutation signature was detected by using SomaticSignatures^54^ v2.20.0. DELLY^55^ v0.8.1, BreakDancer^56^ v1.3.6 and CREST^57^ v2016.12.07 were used to identify the genomic translocations. The mutations results of malignant diseases were downloaded from TCGA studies (https://portal.gdc.cancer.gov/). The mutations results of hepatocellular adenomas (HA) and breast fibroadenomas (BFT) were derived from published studies^24, 25, 26^.

### Copy number analysis

DNA from BPH and matched control samples from blood tissue were analyzed by Affymetrix SNP 6.0 arrays to detect the regions of somatic copy number alteration. Quality control, segmentation and copy number analysis were performed as previously described^23^. The copy number alterations of malignant diseases were downloaded from TCGA portal (https://portal.gdc.cancer.gov/). Segments with log_2_-ratio >0.3 were defined as genomic amplifications, and log_2_-ratio <-0.3 were defined as genomic deletions.

### RNA-seq platform, data procession and analysis pipeline

RNA-seq library for BPH and control samples from patients without BPH symptoms were generated using Poly-A and Ribo-Zero kits. RNA-seq was performed in the Genomics Core from Weill Cornell Medicine under standard protocol and pipeline for 75bp paired-end sequencing. Reads were mapped to the human reference genome sequence (hg19) using STAR^58^ v2.4.0j. Then the resulting BAM files were subsequently converted into mapped-read format (MRF) using RSEQtools^59^ v0.6. The read count of each gene was calculated via HTSeq^60^ v0.11.1 using GENCODE as reference gene annotation set. Quantification of gene expression was performed via RSEQtools^59^ v0.6, and expression levels (RPKM) were estimated by counting all nucleotides mapped to each gene and were normalized by the total number of mapped nucleotides (per million) and the gene length (per kb). Fusion genes were detected via FusionSeq^61^ v0.1.2. Combat^62^ v3.20.0 was used to remove the batch effect of different RNA-seq libraries for the downstream gene expression analysis. Heatmap and hierarchical clustering were performed via using correlation distance and Ward’s method. GSEA^33^ v3.0 was performed using JAVA program and run in pre-ranked mode to identify enriched signatures. The GSEA plot, normalized enrichment score and q-values were derived from GSEA output for hallmark signature, and the metabolism related signatures were derived from MSigDB^63^ v6.2 database. Differential expression analysis was performed using the Wilcoxon signed-rank and F-statistic test after transforming the RPKMs via log2(RPKM + 1). Multiple-hypothesis testing was considered by using Benjamini-Hochberg (BH; FDR) correction. The immune score and stromal score were calculated from gene expression of BPH samples via ESTIMATE^64^ v2.0.0. The ImmuneSum was defined as the sum of normalized z-scores from gene expression of immune markers including *PDCD1*, *PDCD1LG2*, *CD274*, *CD8A* and *CD8B*. The comparison of molecular and clinical features between two subgroups was performed using Fisher’s exact test and Wilcoxon signed-rank test. The compounds for each subgroup were identified by using Connectivity Map (CMAP)^41^ with the top most overexpressed and underexpressed genes as the input, and CMAP score >90 were used to select the nominated compounds.

### Enhanced Reduced Representation Bisulfite Sequencing (ERRBS) platform, data procession and analysis pipeline

Genomic DNA was isolated from BPH and control samples, and submitted to the Epigenomics Core of Weill Cornell Medicine under standard protocol and pipeline. The Epigenomics Core facility in Weill Cornell Medicine supported alignment and methylation extraction for ERRBS data^65^. Differentially methylated CpGs (DMCs) were identified by methylKit^66^ and RRBSeeqer^67^ (false discovery rate =5%, and methylation difference more than 10%). Differentially methylated regions (DMRs) were defined as regions containing at least five DMCs within 250bp window. Genomic regions for CpGs were defined according to the following definitions. CGIs (CpG islands) were defined using annotations from RefSeq. CGI shores were defined as the regions encompassing 1kb upstream and downstream of known CGIs. Non-CGIs were defined as regions at least 10kb away from known CGIs. Promoters were defined as the regions encompassing 2kb upstream and downstream of the TSS (transcription start site) of RefSeq genes. Promoter methylation for each gene was calculated by averaging the methylation levels of all CpGs covered in the promoter.

### Effect of *mTOR* inhibition on prostate size

We searched our electronic medical record to identify all adult male patients who received therapy with an *mTOR* inhibitor (Sirolimus, Everolimus, or Temsirolimus) using our institutional i2b2 search tool (IRB 1510016681R003) (Figure S9). Records were manually reviewed in order to identify individuals with CT imaging containing the pelvis before and after therapy. The most proximate CT scan prior to the initiation of therapy and the CT scan as close to 6 months after the initiation of therapy were used. This interval was chosen based on the known time course of prostate size changes in response to finasteride^68^. As negative controls, we identified 12 kidney stone patients over age 35 at the time of baseline CT who had serial CT imagining including the pelvis who did not take 5-alpha-reductase inhibitors, have prostate cancer, recurrent urinary tract infections, or a history of prostatic surgery. In order to establish a signal window, we performed an initial unblinded pilot including patients who underwent androgen deprivation therapy as a positive control and kidney stone patients as a negative control. We determined that a decrease in prostate size of >12.5% in sequential CT scans would have captured 9/10 androgen deprivation therapy patients from CT scans spaced ∼6 months apart, and would be >2 standard deviations from the mean decrease in prostate size of the kidney stone patients.

We then extracted accession numbers from both kidney stone and *mTOR* inhibitor patients and using a random number generator placed them in arbitrary order for review. A radiologist then reviewed these scans by using accession numbers unaware of treatment assignment (*mTOR* inhibitor or kidney stone) or whether it was a baseline or follow up study. The prostate was measured in two dimensions in the axial slice with the greatest apparent prostate area, with area computed as anterior-posterior x transverse measurements. When unclear, the prostate was measured in two axial slices and the maximum area utilized.

Following review, scans were then re-identified, and subjects with a baseline axial prostate size <1000 mm^2^ were excluded from further analysis. Subjects where the blinded review showed a >12.5% decrease in area, defined as initial area-follow up area)/ initial area, and then underwent a subsequent blinded review by an urologist (JS). Agreement was necessary between both reviews for a decrease to be considered true: when urology review did not identify a decrease >12.5% and this differed from radiology review by <20% urology review was prioritized. For subjects where both reviewers agreed on the decrease in area, initial radiology review dimensions were utilized. When there was a >20% discrepancy in measurements (irrespective of degree), scans were re-reviewed blinded by radiology, blinded, and these measurements utilized.

## Data Availability

The dbGap submission of WGS and WES data is in process. The SNP array data has been deposited in GEO under the accession GSE124187, RNA-seq data has been deposited in GEO under the accession GSE132714, and ERRBS data has been deposited in GEO under the accession GSE123111.

## Author contributions

C.E.B., A.S. and D.L. designed research studies. D.L., J.S., R.S.G., D.R., A.V., D.P., V.R., J.F., H.P., D.L., D.T., K.S., Z.W., A.T., R.L., B.C., A.F.O. and J.M.M. conducted experiments and acquired data. D.L., J.S., C.E.B. and A.S. analyzed the data. D.L., A.S. and C.E.B. wrote the manuscript. J.S., A.R., A.T., R.L., B.C., A.F.O., J.M.M., F.D., O.E., M.A.R., helped to revise the manuscript.

## Acknowledgement

We are grateful to the benign prostatic hyperplasia patients and families who contributed to this research. We thank the WCM Genomics and Epigenomics Core Facility, and all the surgeons, pathologists, research coordinators, and trainees to contributed to patient enrollment and tissue collection. This work was supported by: NCATS (CTCS: UL1 RR 024996), a Urology Care Foundation Rising Star in Urology Research Award (C.E.B.), Damon Runyon Cancer Research Foundation MetLife Foundation Family Clinical Investigator Award (C.E.B.), the Prostate Cancer Foundation (C.E.B), the Prostate Cancer Foundation Young Investigator Award (D.L), the Frederick J. and Theresa Dow Wallace Fund of the New York Community Trust (J.S.), and Damon Runyon Cancer Research Foundation Physician Scientist Training Award (J.S.).

## Reference

1. Bushman W. Etiology, epidemiology, and natural history of benign prostatic hyperplasia. The Urologic clinics of North America 36, 403–415, v (2009).

2. Vuichoud C, Loughlin KR. Benign prostatic hyperplasia: epidemiology, economics and evaluation. The Canadian journal of urology 22 **Suppl 1**, 1–6 (2015).

3. Chughtai B, et al. Benign prostatic hyperplasia. Nature reviews Disease primers 2, 16031 (2016).

4. Lim KB. Epidemiology of clinical benign prostatic hyperplasia. Asian journal of urology 4, 148–151 (2017).

5. Calogero AE, Burgio G, Condorelli RA, Cannarella R, La Vignera S. Epidemiology and risk factors of lower urinary tract symptoms/benign prostatic hyperplasia and erectile dysfunction. The aging male : the official journal of the International Society for the Study of the Aging Male, 1–8 (2018).

6. McVary KT, et al. Update on AUA guideline on the management of benign prostatic hyperplasia. The Journal of urology 185, 1793–1803 (2011).

7. Skinder D, Zacharia I, Studin J, Covino J. Benign prostatic hyperplasia: A clinical review. JAAPA : official journal of the American Academy of Physician Assistants 29, 19–23 (2016).

8. Sarma AV, Wei JT. Clinical practice. Benign prostatic hyperplasia and lower urinary tract symptoms. The New England journal of medicine 367, 248–257 (2012).

9. Bechis SK, Otsetov AG, Ge R, Olumi AF. Personalized medicine for the management of benign prostatic hyperplasia. The Journal of urology 192, 16–23 (2014).

10. Van Asseldonk B, Barkin J, Elterman DS. Medical therapy for benign prostatic hyperplasia: a review. The Canadian journal of urology 22 **Suppl 1**, 7–17 (2015).

11. Nair SM, Pimentel MA, Gilling PJ. Evolving and investigational therapies for benign prostatic hyperplasia. The Canadian journal of urology 22 **Suppl 1**, 82–87 (2015).

12. Negri E, et al. Family history of cancer and the risk of prostate cancer and benign prostatic hyperplasia. International journal of cancer 114, 648–652 (2005).

13. Loeb S, Kettermann A, Carter HB, Ferrucci L, Metter EJ, Walsh PC. Prostate volume changes over time: results from the Baltimore Longitudinal Study of Aging. The Journal of urology 182, 1458–1462 (2009).

14. Wang Z, Olumi AF. Diabetes, growth hormone-insulin-like growth factor pathways and association to benign prostatic hyperplasia. Differentiation; research in biological diversity 82, 261–271 (2011).

15. Wang S, et al. Body mass index and risk of BPH: a meta-analysis. Prostate cancer and prostatic diseases 15, 265–272 (2012).

16. Dhanasekaran SM, et al. Molecular profiling of human prostate tissues: insights into gene expression patterns of prostate development during puberty. FASEB journal : official publication of the Federation of American Societies for Experimental Biology 19, 243–245 (2005).

17. Descazeaud A, et al. BPH gene expression profile associated to prostate gland volume. Diagnostic molecular pathology : the American journal of surgical pathology, part B 17, 207–213 (2008).

18. Winchester D, et al. SPINK1 Promoter Variants Are Associated with Prostate Cancer Predisposing Alterations in Benign Prostatic Hyperplasia Patients. Anticancer research 35, 3811–3819 (2015).

19. Song L, Shen W, Zhang H, Wang Q, Wang Y, Zhou Z. Differential expression of androgen, estrogen, and progesterone receptors in benign prostatic hyperplasia. Bosnian journal of basic medical sciences 16, 201–208 (2016).

20. Wang Z, et al. Androgenic to oestrogenic switch in the human adult prostate gland is regulated by epigenetic silencing of steroid 5alpha-reductase 2. The Journal of pathology 243, 457–467 (2017).

21. Middleton LW, et al. Genomic analysis of benign prostatic hyperplasia implicates cellular re-landscaping in disease pathogenesis. JCI insight 5, (2019).

22. Barbieri CE, et al. Exome sequencing identifies recurrent SPOP, FOXA1 and MED12 mutations in prostate cancer. Nature genetics 44, 685–689 (2012).

23. Cancer Genome Atlas Research N. The Molecular Taxonomy of Primary Prostate Cancer. Cell 163, 1011–1025 (2015).

24. Lim WK, et al. Exome sequencing identifies highly recurrent MED12 somatic mutations in breast fibroadenoma. Nature genetics 46, 877–880 (2014).

25. Tan J, et al. Genomic landscapes of breast fibroepithelial tumors. Nature genetics 47, 1341–1345 (2015).

26. Pilati C, et al. Genomic profiling of hepatocellular adenomas reveals recurrent FRK-activating mutations and the mechanisms of malignant transformation. Cancer cell 25, 428–441 (2014).

27. Nickel JC. Benign prostatic hyperplasia: does prostate size matter? Reviews in urology 5 **Suppl 4**, S12–17 (2003).

28. Al-Khalil S, Ibilibor C, Cammack JT, de Riese W. Association of prostate volume with incidence and aggressiveness of prostate cancer. Research and reports in urology 8, 201–205 (2016).

29. Alexandrov LB, et al. Signatures of mutational processes in human cancer. Nature 500, 415–421 (2013).

30. Pfeifer GP. Mutagenesis at methylated CpG sequences. Current topics in microbiology and immunology 301, 259–281 (2006).

31. Baca SC, et al. Punctuated evolution of prostate cancer genomes. Cell 153, 666–677 (2013).

32. Zhao H, Ramos CF, Brooks JD, Peehl DM. Distinctive gene expression of prostatic stromal cells cultured from diseased versus normal tissues. Journal of cellular physiology 210, 111–121 (2007).

33. Subramanian A, et al. Gene set enrichment analysis: a knowledge-based approach for interpreting genome-wide expression profiles. Proceedings of the National Academy of Sciences of the United States of America 102, 15545–15550 (2005).

34. Fenner A. BPH: Disrupting AR signalling promotes inflammation. Nature reviews Urology 13, 631 (2016).

35. Zhang B, et al. Non-Cell-Autonomous Regulation of Prostate Epithelial Homeostasis by Androgen Receptor. Molecular cell 63, 976–989 (2016).

36. Kohler J, et al. Lestaurtinib inhibits histone phosphorylation and androgen-dependent gene expression in prostate cancer cells. PloS one 7, e34973 (2012).

37. Keszei AF, et al. Structure of an SspH1-PKN1 complex reveals the basis for host substrate recognition and mechanism of activation for a bacterial E3 ubiquitin ligase. Molecular and cellular biology 34, 362–373 (2014).

38. Tomlins SA, et al. Integrative molecular concept modeling of prostate cancer progression. Nature genetics 39, 41–51 (2007).

39. Henry GH, et al. A Cellular Anatomy of the Normal Adult Human Prostate and Prostatic Urethra. Cell reports 25, 3530–3542 e3535 (2018).

40. Lamb J, et al. The Connectivity Map: using gene-expression signatures to connect small molecules, genes, and disease. Science 313, 1929–1935 (2006).

41. Subramanian A, et al. A Next Generation Connectivity Map: L1000 Platform and the First 1,000,000 Profiles. Cell 171, 1437–1452 e1417 (2017).

42. Shi YF, et al. TRAF6 regulates proliferation of stromal cells in the transition and peripheral zones of benign prostatic hyperplasia via Akt/mTOR signaling. The Prostate 78, 193–201 (2018).

43. Zhang N, et al. MicroRNA expression profiles in benign prostatic hyperplasia. Molecular medicine reports 17, 3853–3858 (2018).

44. Lesovaya EA, et al. Rapatar, a nanoformulation of rapamycin, decreases chemically-induced benign prostate hyperplasia in rats. Oncotarget 6, 9718–9727 (2015).

45. Mehine M, et al. Characterization of uterine leiomyomas by whole-genome sequencing. The New England journal of medicine 369, 43–53 (2013).

46. Ehrlich M. DNA methylation in cancer: too much, but also too little. Oncogene 21, 5400–5413 (2002).

47. Jiang L, et al. Global hypomethylation of genomic DNA in cancer-associated myofibroblasts. Cancer research 68, 9900–9908 (2008).

48. Torano EG, Petrus S, Fernandez AF, Fraga MF. Global DNA hypomethylation in cancer: review of validated methods and clinical significance. Clinical chemistry and laboratory medicine 50, 1733–1742 (2012).

49. Li H, Durbin R. Fast and accurate short read alignment with Burrows-Wheeler transform. Bioinformatics 25, 1754–1760 (2009).

50. McKenna A, et al. The Genome Analysis Toolkit: a MapReduce framework for analyzing next-generation DNA sequencing data. Genome research 20, 1297–1303 (2010).

51. Cibulskis K, et al. Sensitive detection of somatic point mutations in impure and heterogeneous cancer samples. Nature biotechnology 31, 213–219 (2013).

52. Koboldt DC, et al. VarScan 2: somatic mutation and copy number alteration discovery in cancer by exome sequencing. Genome research 22, 568–576 (2012).

53. Wang K, Li M, Hakonarson H. ANNOVAR: functional annotation of genetic variants from high-throughput sequencing data. Nucleic acids research 38, e164 (2010).

54. Gehring JS, Fischer B, Lawrence M, Huber W. SomaticSignatures: inferring mutational signatures from single-nucleotide variants. Bioinformatics 31, 3673–3675 (2015).

55. Rausch T, Zichner T, Schlattl A, Stutz AM, Benes V, Korbel JO. DELLY: structural variant discovery by integrated paired-end and split-read analysis. Bioinformatics 28, i333–i339 (2012).

56. Fan X, Abbott TE, Larson D, Chen K. BreakDancer: Identification of Genomic Structural Variation from Paired-End Read Mapping. Current protocols in bioinformatics 45, 15 16 11-11 (2014).

57. Wang J, et al. CREST maps somatic structural variation in cancer genomes with base-pair resolution. Nature methods 8, 652–654 (2011).

58. Dobin A, et al. STAR: ultrafast universal RNA-seq aligner. Bioinformatics 29, 15–21 (2013).

59. Habegger L, et al. RSEQtools: a modular framework to analyze RNA-Seq data using compact, anonymized data summaries. Bioinformatics 27, 281–283 (2011).

60. Anders S, Pyl PT, Huber W. HTSeq--a Python framework to work with high-throughput sequencing data. Bioinformatics 31, 166–169 (2015).

61. Sboner A, et al. FusionSeq: a modular framework for finding gene fusions by analyzing paired-end RNA-sequencing data. Genome biology 11, R104 (2010).

62. Johnson WE, Li C, Rabinovic A. Adjusting batch effects in microarray expression data using empirical Bayes methods. Biostatistics 8, 118–127 (2007).

63. Liberzon A, Subramanian A, Pinchback R, Thorvaldsdottir H, Tamayo P, Mesirov JP. Molecular signatures database (MSigDB) 3.0. Bioinformatics 27, 1739–1740 (2011).

64. Yoshihara K, et al. Inferring tumour purity and stromal and immune cell admixture from expression data. Nature communications 4, 2612 (2013).

65. Garrett-Bakelman FE, et al. Enhanced reduced representation bisulfite sequencing for assessment of DNA methylation at base pair resolution. Journal of visualized experiments : JoVE, e52246 (2015).

66. Akalin A, et al. methylKit: a comprehensive R package for the analysis of genome-wide DNA methylation profiles. Genome biology 13, R87 (2012).

67. Pan H, et al. Epigenomic evolution in diffuse large B-cell lymphomas. Nature communications 6, 6921 (2015).

68. Gormley GJ, et al. The effect of finasteride in men with benign prostatic hyperplasia. The Finasteride Study Group. The New England journal of medicine 327, 1185–1191 (1992).

