## Supplementary Figures for "Integrative genomic, transcriptomic, and epigenomic analyses of benign prostatic hyperplasia reveal new options for therapy"

Figure S1. Circos plots of 18 BPH samples.

Figure S2. BPH transcription signature is not transitional zone specific.

Figure S3. Differentially methylated CpGs and regions found in BPH by ERRBS data.

Figure S4. Hierarchical clustering and heatmap of 18 BPH samples based on prostate stromal signature[1].

Figure S5. Hierarchical clustering and heatmap of GSE101486 samples based on prostate stromal signature[1].

Figure S6. Metabolism dysregulation between two subgroups from both current and GSE101486 studies via GSEA.

Figure S7. Stromal signature enrichment in subgroup BPH-A from single-cell RNA-seq study[2] via GSEA.

Figure S8. Beeswarm plot of stromal enrichment difference between BPH subgroups.

Figure S9. CONSORT diagram detailing work-flow for clinical study.

Figure S10. Boxplot of prostate size changes on patients with and without *mTOR* inhibitors.

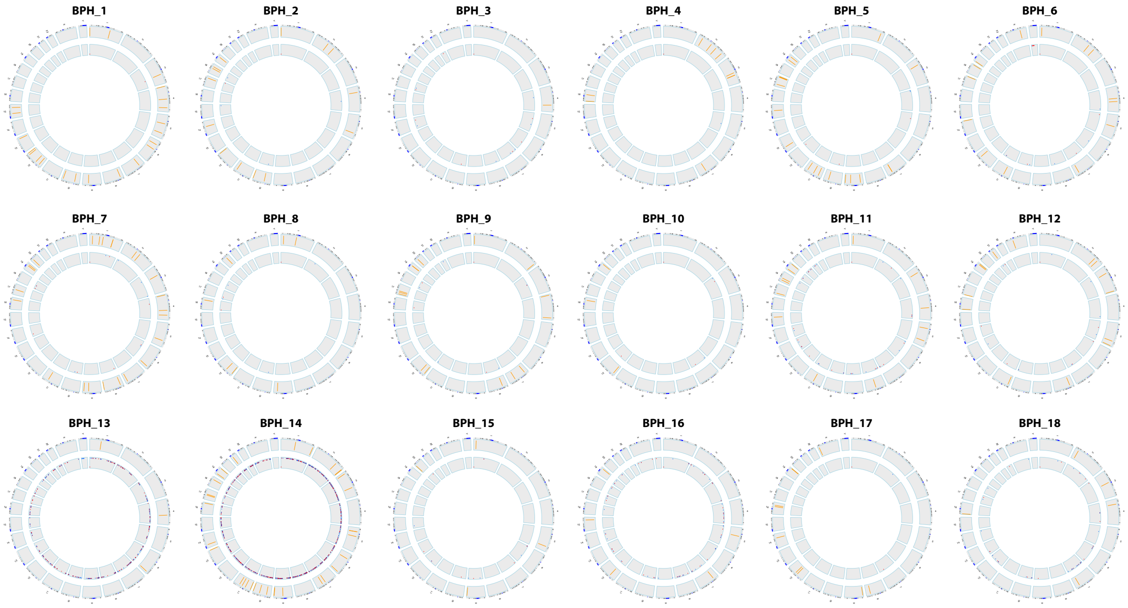


Figure S1. Circos plots of 18 BPH samples. The rings from outer to inner represent somatic coding mutations, copy number alterations and fusion genes respectively.

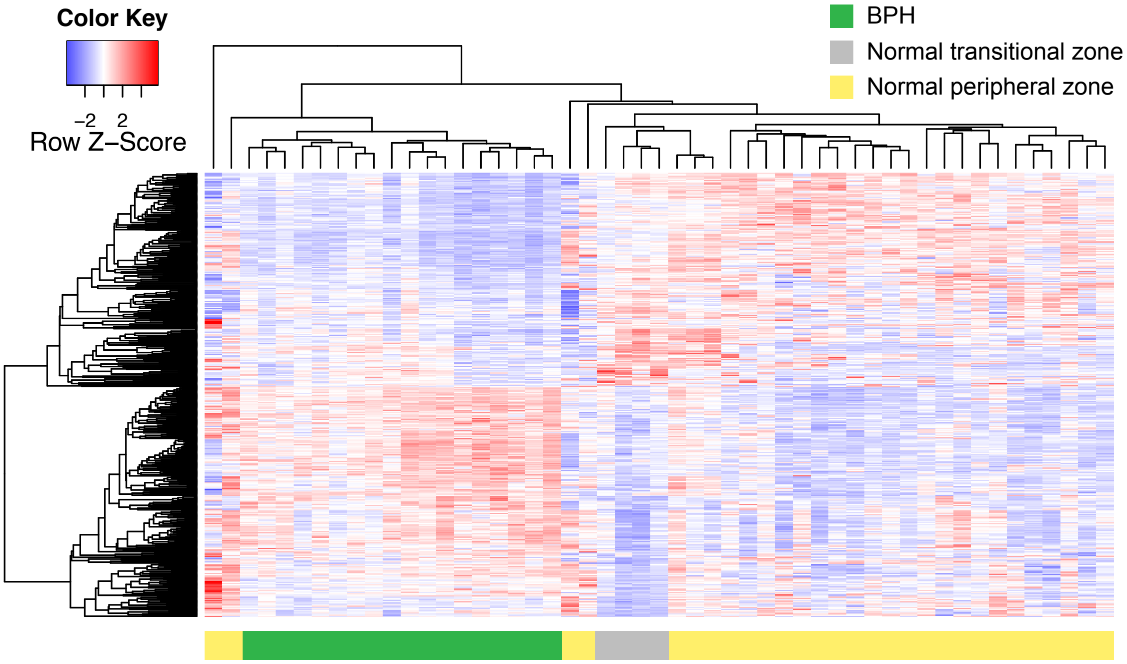


Figure S2. BPH transcription signature is not transitional zone specific. The hierarchical clustering and heatmap of 18 BPH, 4 normal transitional zone, and 29 normal peripheral zone samples, based on BPH transcription signature from Figure 2B.

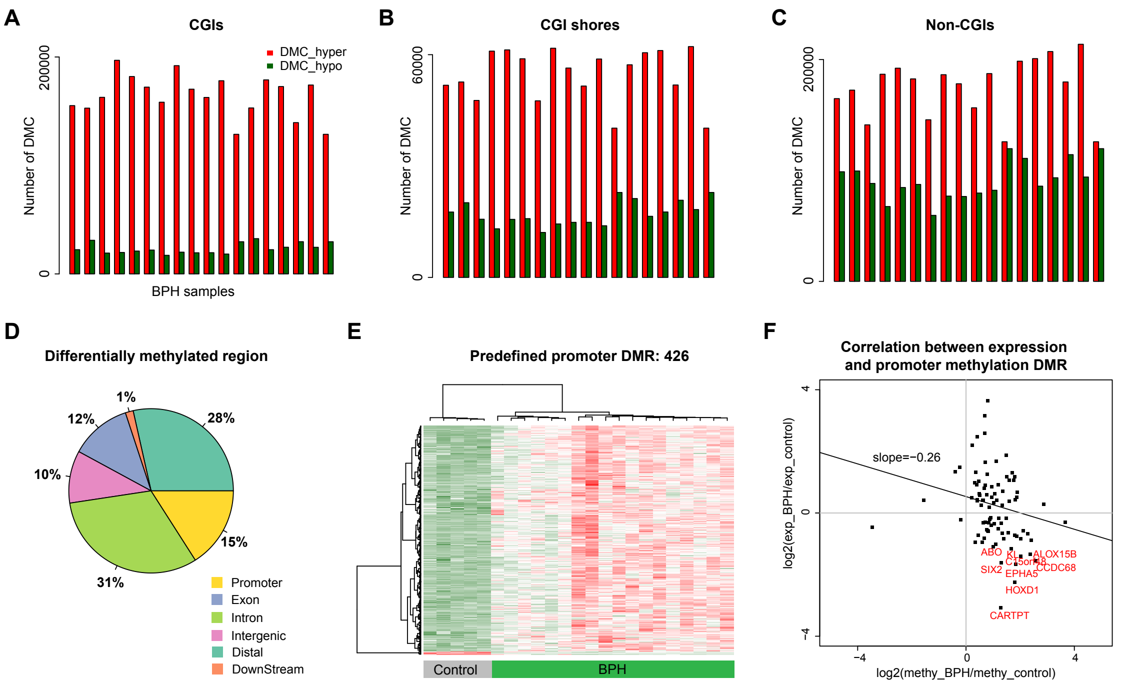


Figure S3. Differentially methylation CpGs and regions found in BPH by ERRBS data.

(A) Hypermethylation domination found in CpG islands (CGIs). Red bar represents the hypermethylated CpGs, and green bar represents the hypomethylated CpGs.

(B) Hypermethylation domination found in CGI shores.

(C) Hypermethylation domination found in non-CGIs.

(D) Pie chart of differentially methylated regions between BPH and control samples among different genomic regions. Different colors denote different genomic related regions

(E) Hierarchical clustering and heatmap of DMRs in promoter between BPH and control samples.

(F) The negative correlation between transcription and methylation signatures, and the examples of epigenetically silent genes are shown in red color.

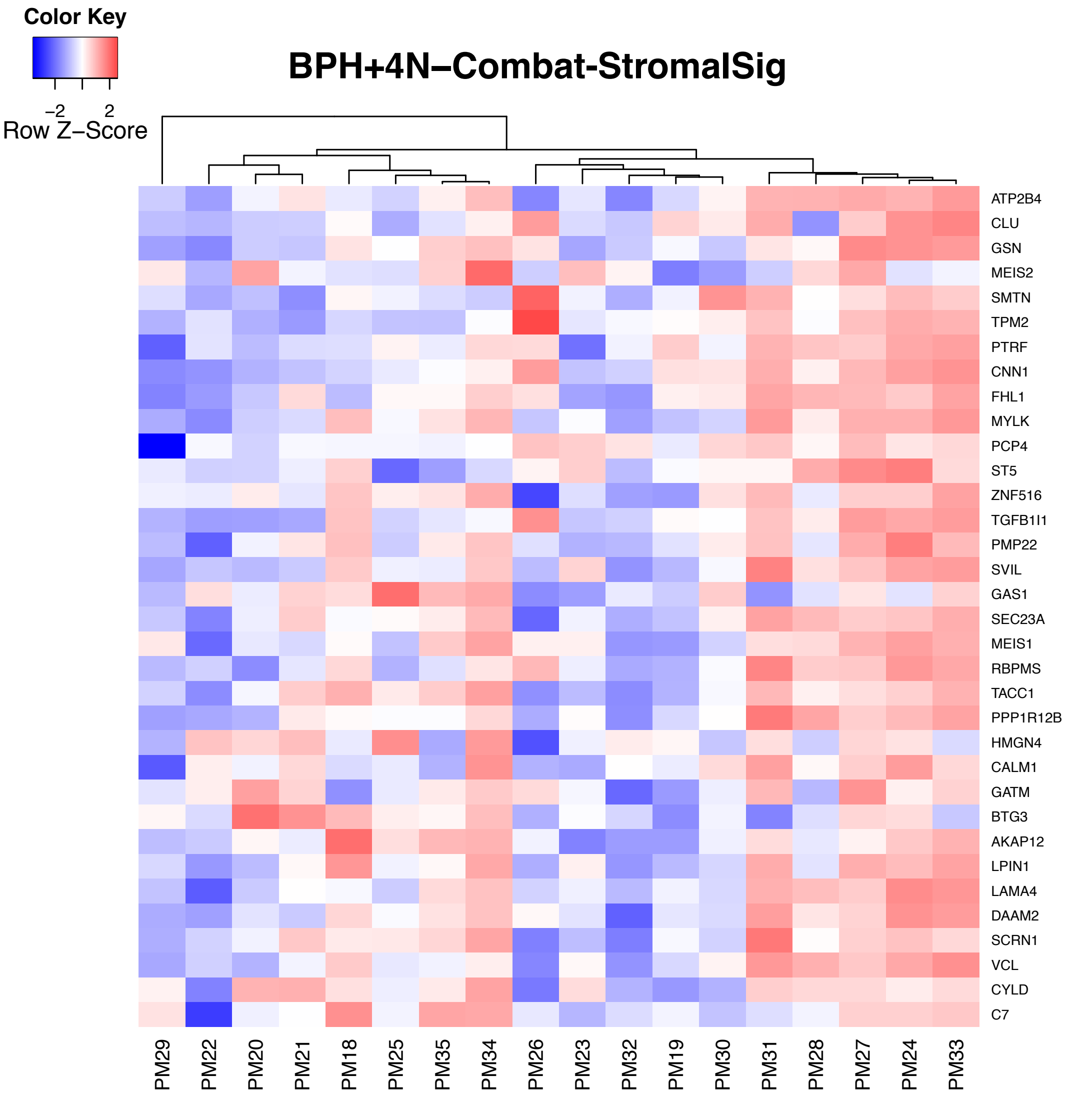


Figure S4. Hierarchical clustering and heatmap of 18 BPH samples based on prostate stromal signature[1].

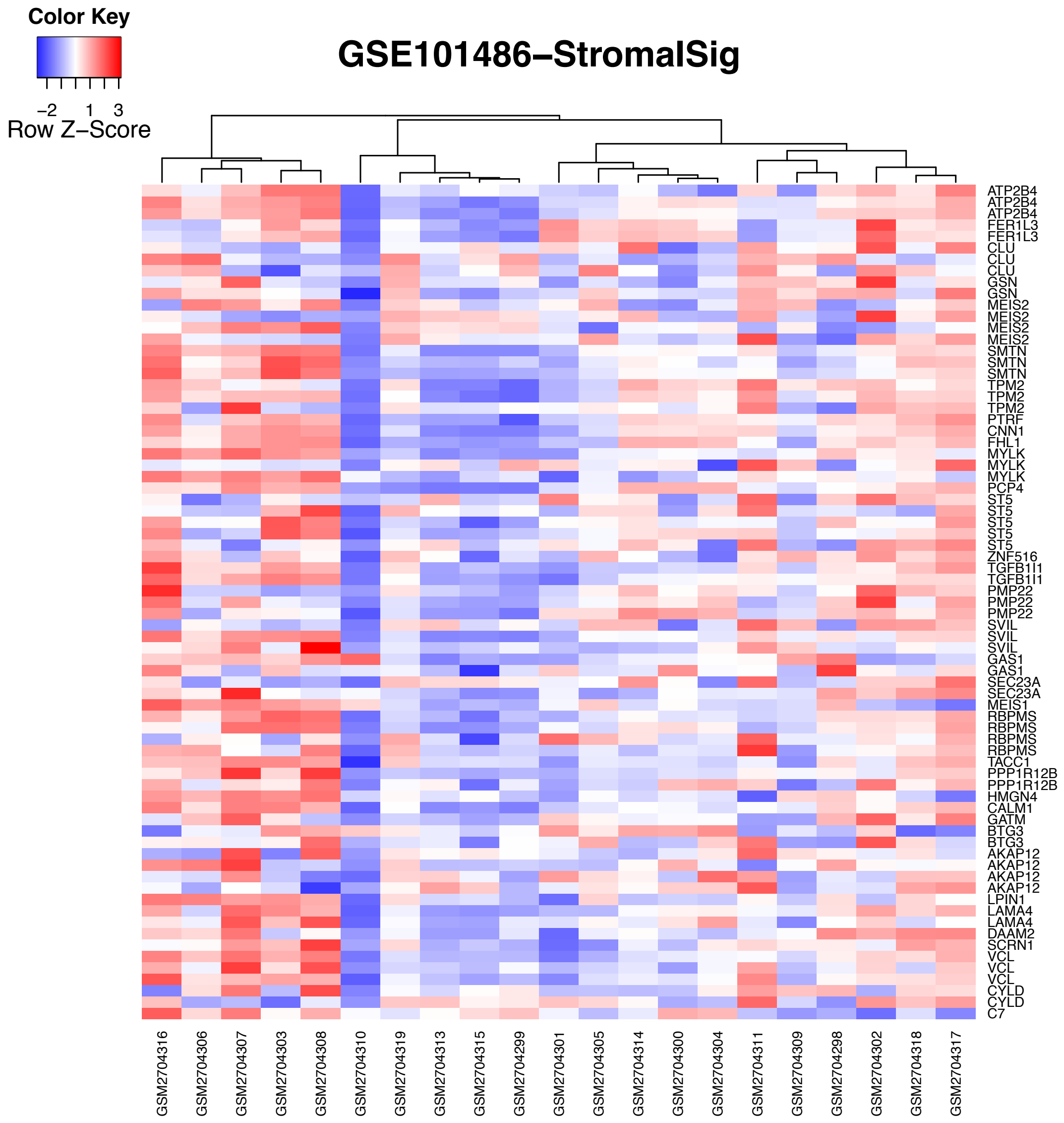


Figure S5. Hierarchical clustering and heatmap of GSE101486 samples based on prostate stromal signature[1].

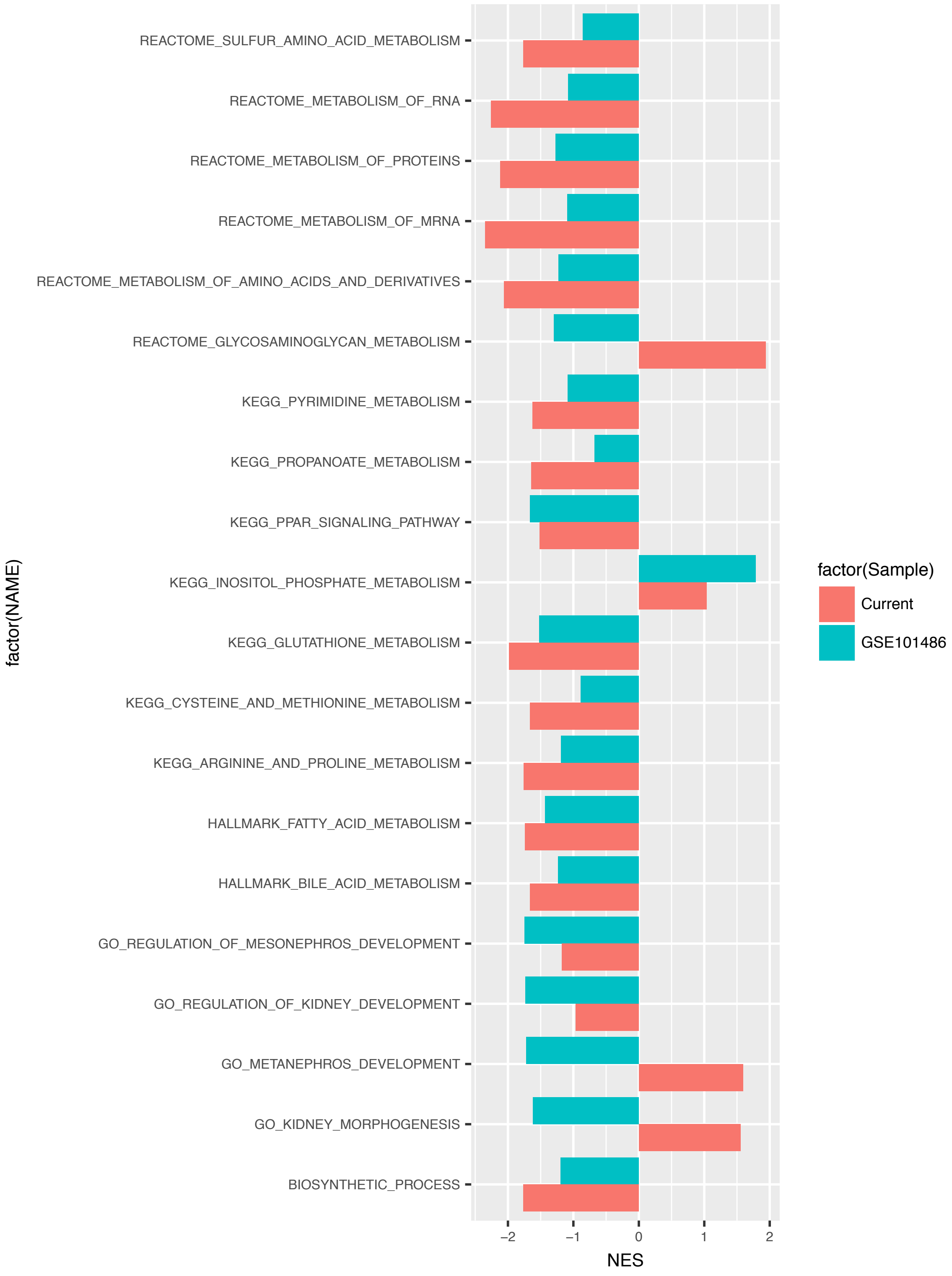


Figure S6. Metabolism dysregulation between two subgroups from both current and GSE101486 studies via GSEA. The x-axis represents the normalized enrichment score, and different colors represent different studies.

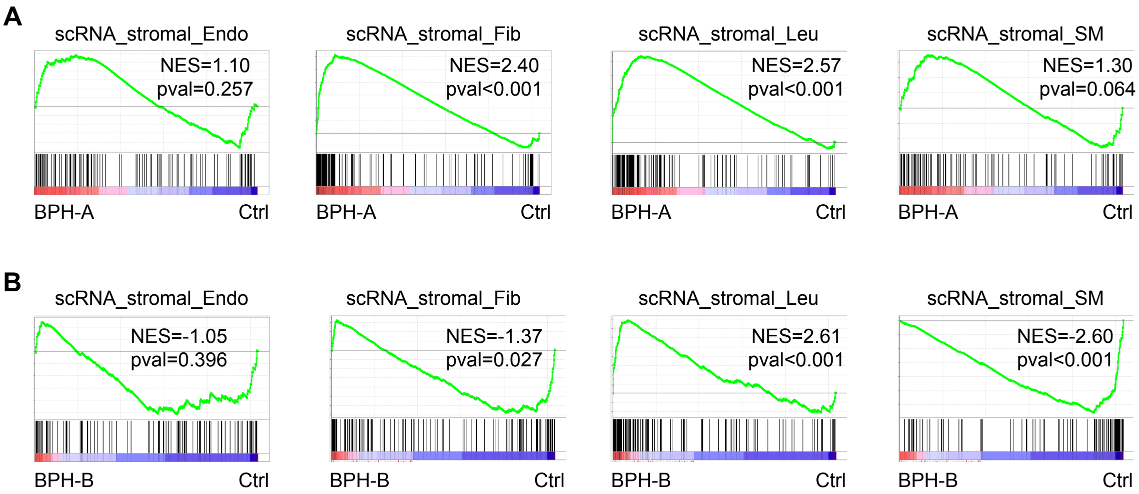


Figure S7. Stromal signature enrichment in subgroup BPH-A from single-cell RNA-seq study[2] via GSEA.

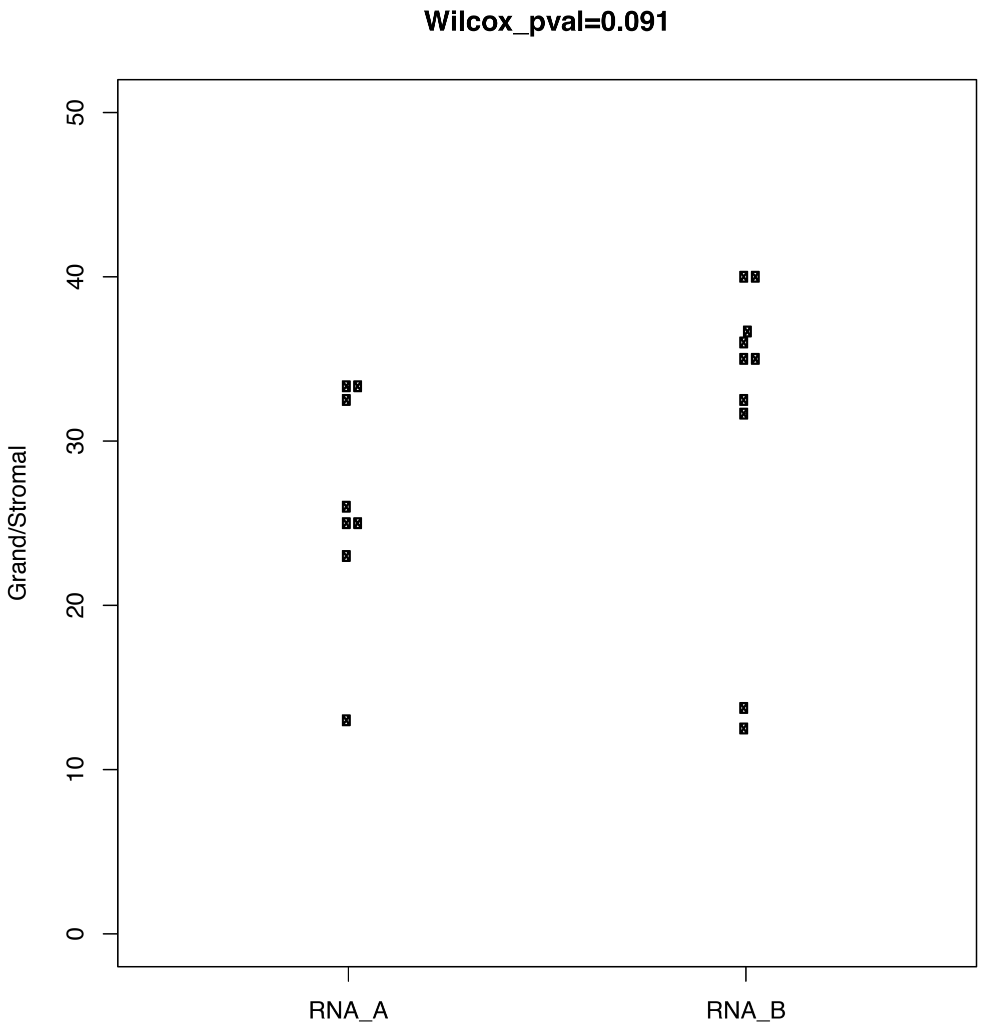


Figure S8. Beeswarm plot of stromal enrichment difference between BPH subgroups. The y-axis denotes the average of grand/stromal cell ratio from multiple slides for each BPH sample. The lower ratio represents higher stromal enrichment. The comparison p-value was calculated from the grand/stromal cell ratio difference between two BPH subgroups via Wilcoxon rank sum test.

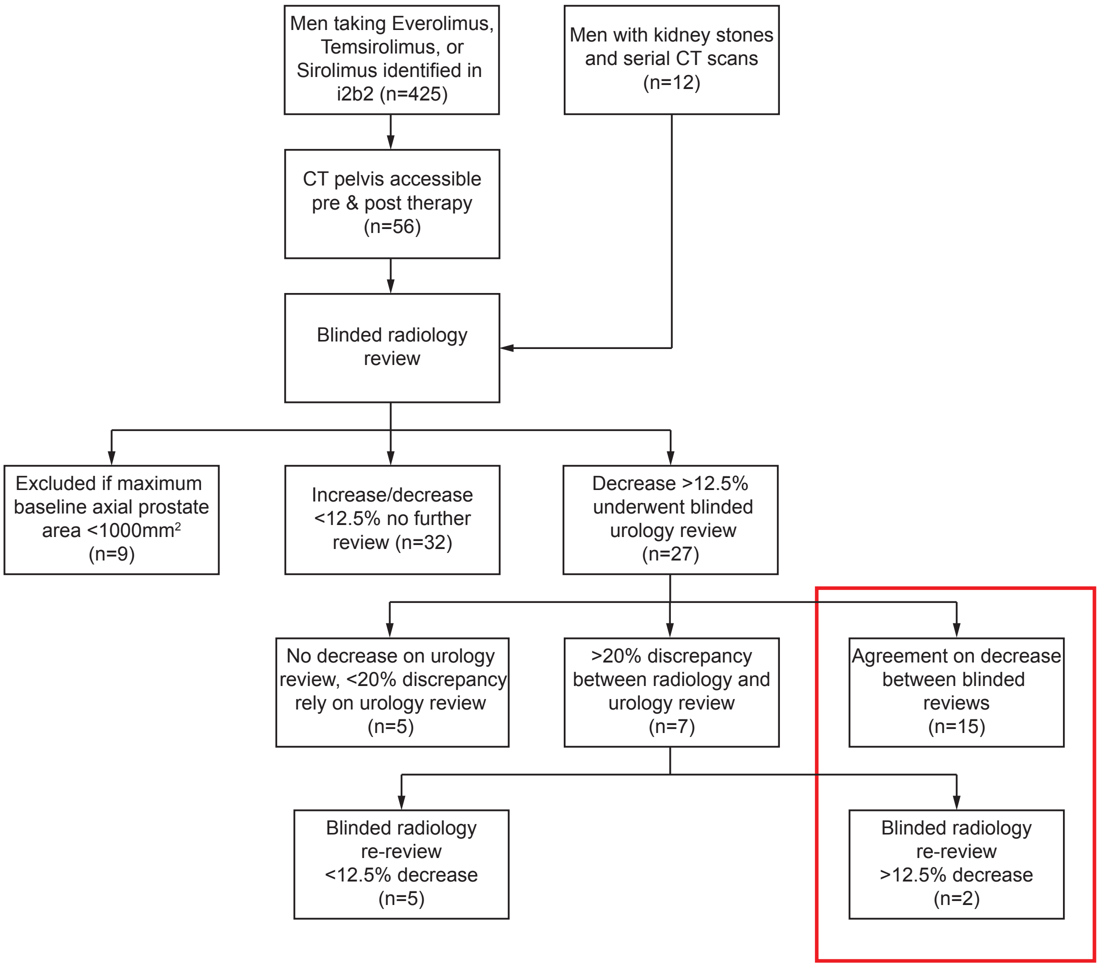


Figure S9. Figure. CONSORT diagram detailing work-flow for clinical study. Box denotes patients considered to have a significant decrease in prostate size.

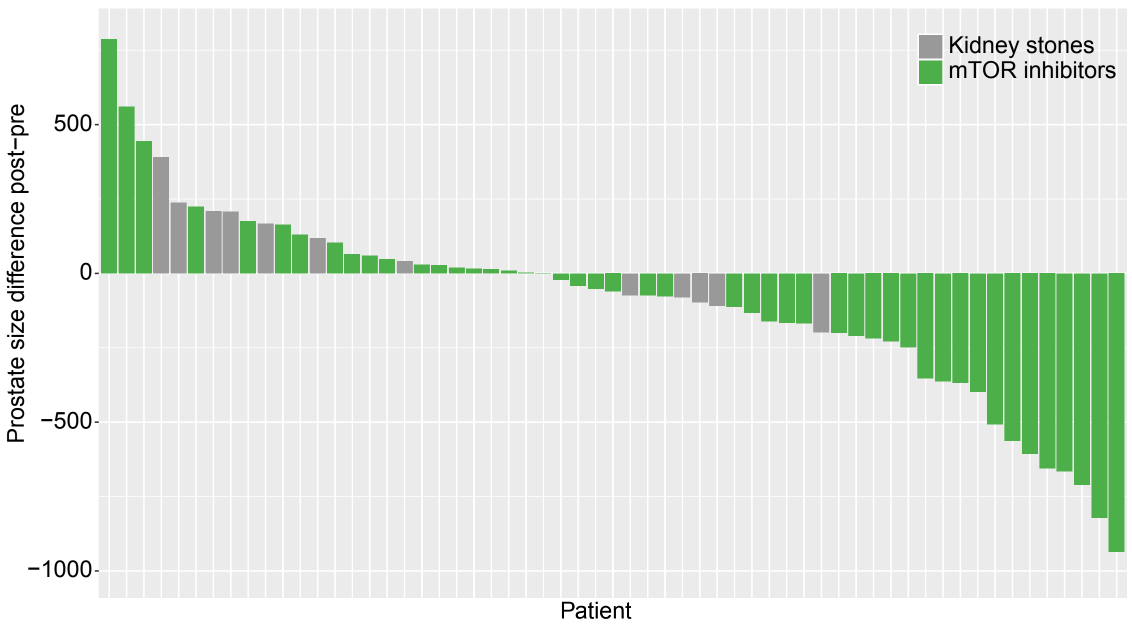


Figure S10. Boxplot of prostate size changes in axial area on patients with and without *mTOR* inhibitors. Different color represents different treatment.

**Reference**

1 Tomlins, S. A. *et al.* Integrative molecular concept modeling of prostate cancer progression. *Nature genetics* **39**, 41-51, doi:10.1038/ng1935 (2007).

2 Henry, G. H. *et al.* A Cellular Anatomy of the Normal Adult Human Prostate and Prostatic Urethra. *Cell reports* **25**, 3530-3542 e3535, doi:10.1016/j.celrep.2018.11.086 (2018).
